# DGAT1-dependent lipid droplet synthesis in microglia attenuates neuroinflammatory responses to lipopolysaccharides

**DOI:** 10.64898/2025.12.18.695188

**Authors:** Frédérick Boisjoly, Danie Majeur, Josephine Louise Robb, Demetra Rodaros, Thierry Alquier

**Affiliations:** Centre de Recherche du Centre Hospitalier de l’Université de Montréal (CRCHUM), Université de Montréal, Montréal, QC, Canada; Departments of Neurosciences, Université de Montréal, Montréal, QC, Canada; Departments of Medicine, Université de Montréal, Montréal, QC, Canada

**Keywords:** Diacylglycerol acyltransferase (DGAT), Lipid Droplet, Microglia, Neuroinflammation

## Abstract

Lipid droplets (LD) are dynamic storage organelles for triglycerides (TG). LD act as a hub that modulates the availability of fatty acids to sustain metabolic needs and the generation of fatty-acid derived signals. Recent evidence demonstrates that LD metabolism regulates immune responses including in microglia, the resident immune cells of the central nervous system. We have previously shown that blocking LD lipolysis in microglia reduces acute pro-inflammatory responses to lipopolysaccharide (LPS) including cytokine and prostanoid synthesis. Here, we investigated the role of diacylglycerol O-acyltransferase 1 (DGAT1), a key enzyme catalyzing the final step of TG synthesis, in microglial LD biogenesis and inflammatory responses induced by LPS. We found that treatment with LPS downregulates specific enzymes in the TG synthesis pathway in primary microglia, including GPAT1, AGPAT3, AGPAT5, and DGAT1, while upregulating TMEM68, a non-canonical TG synthesizing enzyme. Pharmacological inhibition of DGAT1 significantly reduced LD formation in both oleate- and LPS-stimulated conditions, indicating that DGAT1 is essential for inflammation-induced LD synthesis. Moreover, DGAT1 inhibition selectively decreased expression of pro-inflammatory cytokines TNF-α and IL-1β, without affecting IL-6, CCL2, or the anti-inflammatory cytokine TGF-β. These findings show, that DGAT1-dependent LD formation acts as an important modulator of microglial inflammatory signaling.

## INTRODUCTION

Microglia, the main immune cells of the central nervous system, play a crucial role in monitoring, defending, and remodeling the brain parenchyma ^1–4^. These functions require microglia to rapidly adapt their metabolism to sustain their activity ^5,6^. Amongst the pathways that support this flexibility, lipid metabolism has emerged as a key mechanism enabling microglial responses. Recent evidence highlights the role of intracellular triglycerides (TG) storage as an important source of fatty acids acting as metabolic substrates and precursors of lipid signals that sustain microglia functions, but their impact and underlying mechanisms on microglial activation remains poorly understood ^7–9^.

Lipid droplets (LD) are intracellular storage organelles composed of neutral lipids, mainly TG and cholesterol esters ^10^. In microglia, LD accumulate in response to pro-inflammatory stimuli like lipopolysaccharide (LPS) ^11–13^ and their presence has been associated with a dysfunctional state during brain aging and in several neurodegenerative diseases^11,14–16^. We recently showed that microglial LD lipolysis via adipose triglyceride lipase (ATGL), the first enzyme responsible for LD breakdown, is required for the synthesis and release of pro-inflammatory cytokines and prostanoids induced by LPS^12^. Consistent with our findings, studies in primary microglia and human microglial cell lines have reported similar results, showing that inhibition of ATGL attenuates the pro-inflammatory response to LPS ^13,17^. Given that LD lipolysis is a key component of microglial activation, we proposed that LD likely serve as a central hub for lipid inflammatory signaling molecules. This raises the question of whether the esterification arm controlling TG synthesis and LD formation also influences microglial activation.

Recent studies in macrophages and adipocytes showed that diacylglycerol O-acyltransferase (DGAT), the enzyme catalyzing the final step of TG synthesis, plays a critical role in regulating LD accumulation and modulating inflammatory responses ^18–21^. Similar mechanisms were recently discovered in human induced pluripotent stem cell (iPSC)-derived microglia from APOE3/APOE4 mutant cell lines, in which blocking both DGAT1 and 2 isoforms under pro-inflammatory conditions reduces microglial acitvation^22^. However, whether and how the TG synthesis pathway is regulated by inflammatory signals and influences the acute pro-inflammatory response in mouse primary microglia remains unknown. Here we demonstrate that acute LPS-induced LD formation in neonatal primary microglia is mediated by DGAT1 and that its pharmacological inhibition reduces microglial pro-inflammatory responses to LPS.

## RESULTS

### LPS downregulates enzymes of the canonical TG-esterification pathway while upregulating the non-canonical TG synthesizing enzyme TMEM68

We and others have reported that LPS induces LD formation in primary microglial^11–13^. To investigate whether LPS-induced LD synthesis in microglia is associated with changes in the expression of the main glycerolipid esterification enzymes (**Fig. 1a**), their expression was measured by RT-qPCR in neonatal primary microglia cultures (mixed sex) treated for 6 h with 0.1 µg/mL of LPS (**Fig. 1b**). Cell viability and cytotoxicity were not affected by LPS after 6 h (**Fig.S1a-b**). Surprisingly, LPS led to a significant reduction of enzymes involved in each step of TG synthesis: GPAT1 expression was reduced by 70%, AGPAT3 by 35%, AGPAT5 by 39% and DGAT1 by 29% (**Fig.1b**). AGPAT1, AGPAT2, AGPAT6 expression was not affected by LPS (**Fig.1b**). Other enzymes of the esterification pathway could not be quantified (GPAT2 and DGAT2), as their expression was below the detection threshold. These results suggest that acute inflammatory microglial activation by LPS selectively downregulates steps of the esterification pathway.

**Figure 1.**
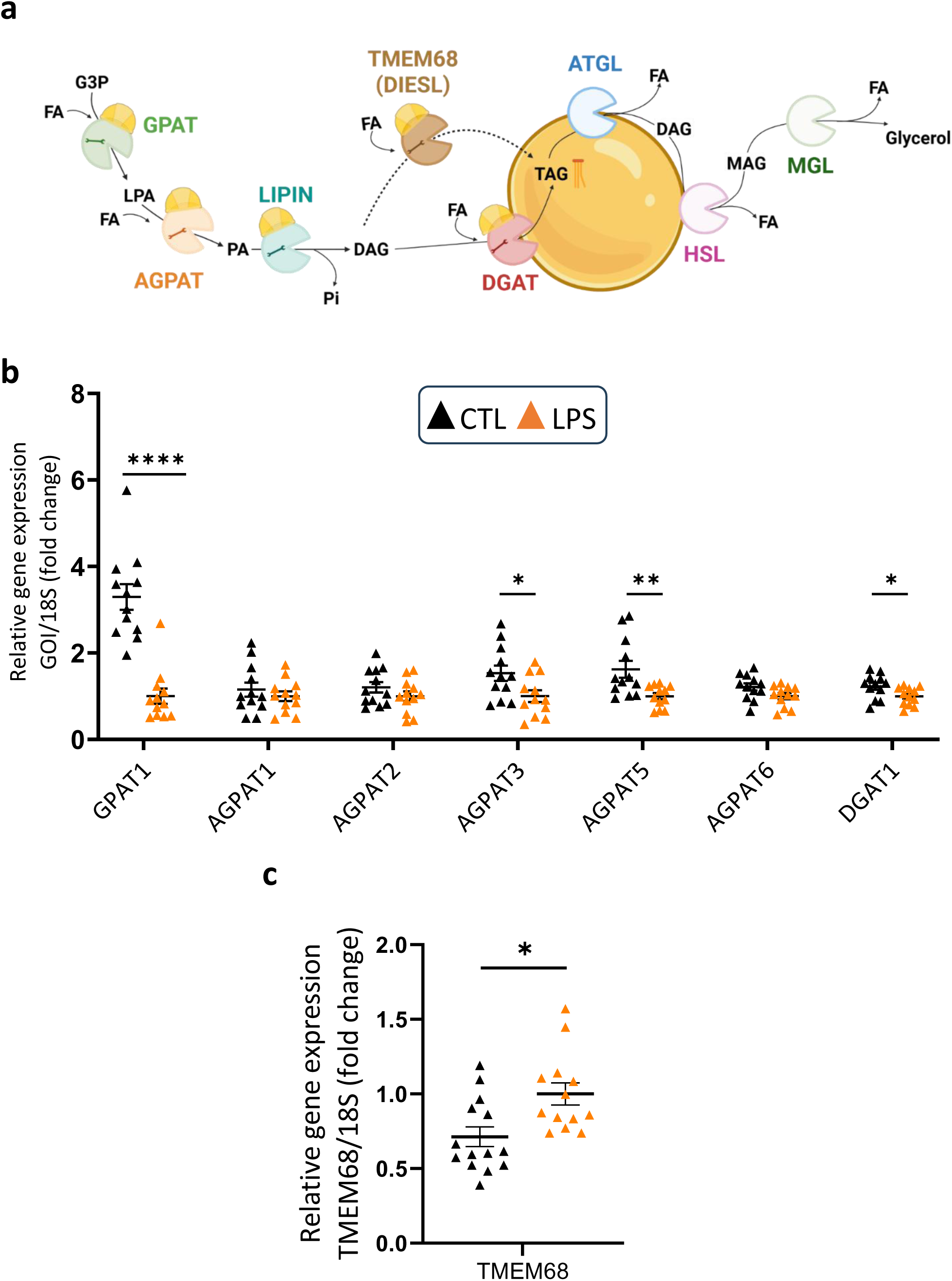
LPS reduces TG-esterification enzymes and increases an alternative TG-synthesis pathway in primary microglia. Primary microglia cultures were treated for 6 h with 0.1 µg/mL LPS. **a)** Scheme illustrating LD formation through alternative TMEM68 (DIESL) and classical TG synthesis enzymes GPAT, AGPAT, LIPIN, and DGAT and LD lipolysis mediated by ATGL, HSL and MGL. **b)** Relative gene expression of enzymes from the classical TG synthesis pathway or was measured by RT-qPCR (n = 12). **c)** Relative gene expression of TMEM68 was measured by RT-qPCR (n = 14). Samples were analyzed by Student’s t-test or Mann–Whitney test. * p < 0.05; ** p < 0.01; **** p < 0.0001. G3P : glycerol-3-phosphate; FA : fatty acids, GPAT : glycerol-3-phosphate acyl-transferase; LPA : lysophosphatidic acid; AGPAT: acylglycerol-3-phosphate acyl-transferase ; PA : phosphatidic acid ; Pi : Phosphate ; DAG : diacylglycerol ; DGAT : diacylglycerol acyl-transferase; TMEM68; Transmembrane protein 68, DIESL; DGAT1/2-independent enzyme synthesizing storage lipids ; TAG : triglycerides ; ATGL : adipose triglyceride lipase ; HSL : hormone-sensitive lipase; MAG : monacylglycerol ; MGL : monoacylglycerol lipase.

Given this coordinated downregulation of the canonical TG esterification pathway, we next asked if the alternative TG-synthesis pathway is modulated by LPS. Recently, TMEM68 (DIESL) has been identified as a novel enzyme capable of generating TG and promoting LD formation independently of the DGAT1/2 pathway^23–25^ (**Fig.1a)**. A recent study showed that brain TG levels are reduced in whole body TMEM68 knockout mice^26^. Furthermore, *Manceau et al.* demonstrated that deletion of TMEM68 in *Drosophila* neurons completely abolishes LD formation^27^. In cultured microglia, we found that TMEM68 expression was upregulated by 29% in response to LPS compared to control (**Fig. 1c**). This suggests that under acute pro-inflammatory conditions, when DGAT1 expression is downregulated, TMEM68 upregulation may contribute to TG synthesis and LD formation.

### Blocking DGAT1 in primary microglia prevents LD formation in oleate-stimulated conditions

To determine whether DGAT1 is the main enzyme contributing to LD formation in microglia, primary microglia were treated with exogenous oleate during 3 h to promote LD formation (**Fig.2a-d**) with or without the DGAT1 inhibitor A922500. Oleate-treated cells were exposed to increasing doses of A922500 to assess a dose response effect (**Fig.2e**). Cell viability and cytotoxicity were not affected by DGAT1 inhibition (**Fig.S1a-b**). In oleate-treated conditions, LD number was on average 25.2 ± 1.3 per cell. Treatment with increasing concentrations of A922500 reduced this mean to 10.0 ± 0.8 LD per cell at 100 nM, 5.0 ± 0.4 LD per cell at 200 nM, and 2.8 ± 0.3 LD per cell at 500 nM (**Fig.2e**). These results indicate that DGAT1 is the major enzyme responsible for oleate-induced LD formation in microglia. For efficient inhibition of LD formation, the 500 nM concentration was chosen for subsequent experiments.

**Figure 2.**
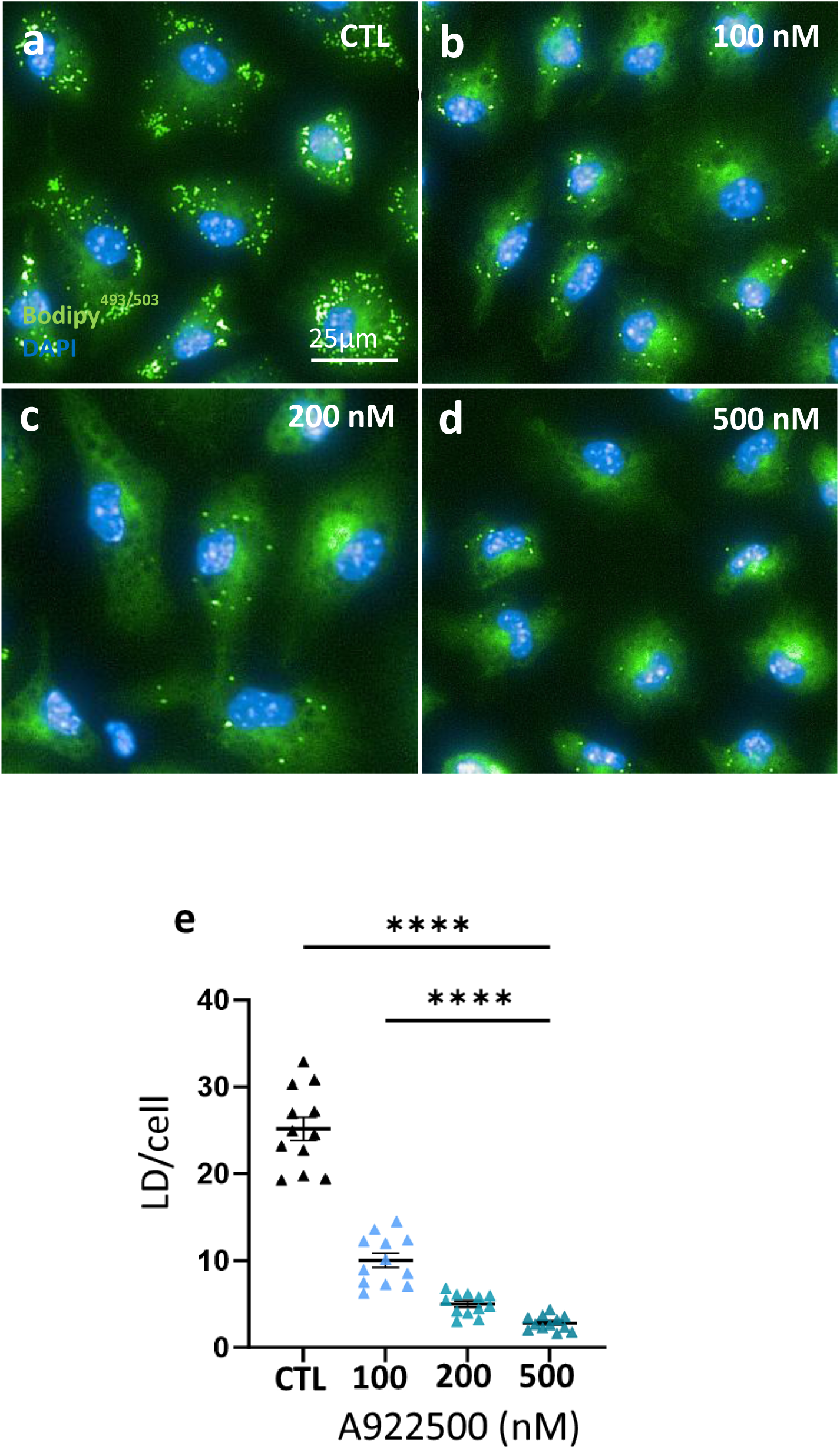
A922500 reduces oleate-induced LD genesis in primary microglia in a dose-dependent manner. Effect of treatment with 0.25 mM oleate pre-complexed with 0.3% BSA, with or without A922500 on LD number. **a-d)** Representative images of primary neonatal microglia pre-treated with oleate (2 h) and maintained (3 h) with oleate and different doses of A922500, stained with BODIPY (green) and Hoechst (blue), Scale bar : 25 µm. **e)** Average number of LD per cell treated with different doses of A922500 (n =12, 9 fields of view/n, ∼ 6400 cells counted/conditions). One-way ANOVA with post-hoc Šidák.

### DGAT1 inhibition suppresses LD formation in LPS-stimulated microglia

We next assessed if DGAT1 inhibition reduces LD formation in response to LPS in the absence of exogenous fatty-acids (**Fig.3a-d**). Primary microglia were pre-treated with A922500 or vehicle (DMSO) for 2 h and subsequently incubated for 6 h with or without LPS in the presence of A922500. Blocking DGAT1 under non-stimulated conditions did not affect either LD number per cell or the proportion of LD-positive cells compared to vehicle (**Fig.3e-f**). As expected, LPS increased LD accumulation from a mean of 1.0 ± 0.2 to 2.5 ± 0.3 LD per cell and raised the proportion of LD-positive cells from 44% to 69% (**Fig.3e-f**). Cell viability and cytotoxicity were not affected by the combination of A922500 (500 nM) and LPS (**Fig.S1a-b**). DGAT1 inhibition significantly reduced LPS-induced LD accumulation to a mean of 1.4 ± 0.2 LD per cell and lowered the percentage of LD-positive cells to 50% (**Fig.3e-f**). Similar results were obtained when primary microglia were incubated with oleate in presence of LPS. As observed in **Fig.2**, DGAT1 inhibition strongly reduced LD number in oleate-treated cells. In presence of oleate, LPS further increased LD number from an average of 15.3 ± 0.5 to a mean of 23.8 ± 0.8 LD per cell (**Fig.S2 a-e**). DGAT1 inhibition strongly reduced LD number to 1.9 ± 0.2 LD per cell in presence of oleate and LPS (**Fig.S2 a-e**). These data imply that while basal LD synthesis appears to be independent of DGAT1 activity, DGAT1 is largely responsible for LPS-induced LD formation with or without added fatty acids.

**Figure 3.**
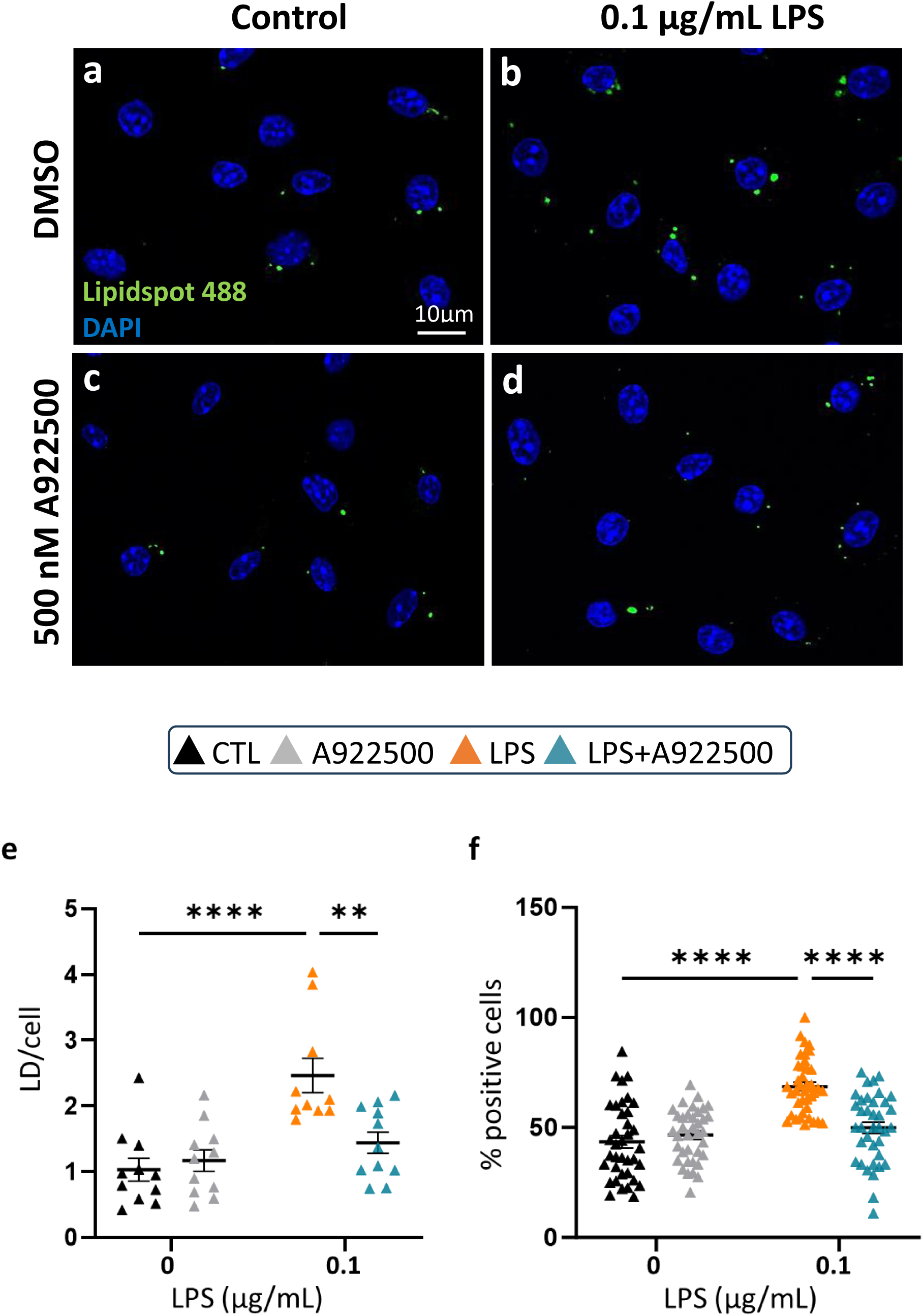
DGAT1 is required for LPS-induced LD synthesis in microglia. Effect of 6 h treatment with 0.1 µg/mL LPS ± 500nM A922500 on LD formation **a-d)** Representative images of primary neonatal microglia during a 6 h treatment with LPS or A922500, stained with Lipidspot 488 (green) and Hoechst (blue), 10 µm scale. **e)** Average number of LD per cell (n = 11-12, 3-4 fields of view/n). **f)** Average number of LD-positive cells (n = 37-40, ∼ 1100 cells counted/conditions). Two-way ANOVA with post-hoc Šídák, **p < 0.01 **** p < 0.0001.

### Pharmacological inhibition of microglial DGAT1 reduces pro-inflammatory cytokine expression

Given that DGAT1 is responsible for microglial LD formation induced by LPS, we next investigated whether its inhibition also affects LPS-induced pro-inflammatory responses. Vehicle or A922500 pre-treated microglia were treated for 6 h with A922500 and/or LPS, and cytokine expression was quantified by RT-qPCR. DGAT1 inhibition reduced LPS-induced expression of TNF-α and IL-1β by 23% and 24% respectively, without significantly affecting IL-6 or CCL2 levels (**Fig.4a**). In addition, expression of the anti-inflammatory cytokine TGF-β remained unchanged following A922500 treatment (**Fig.4a**).

**Figure 4.**
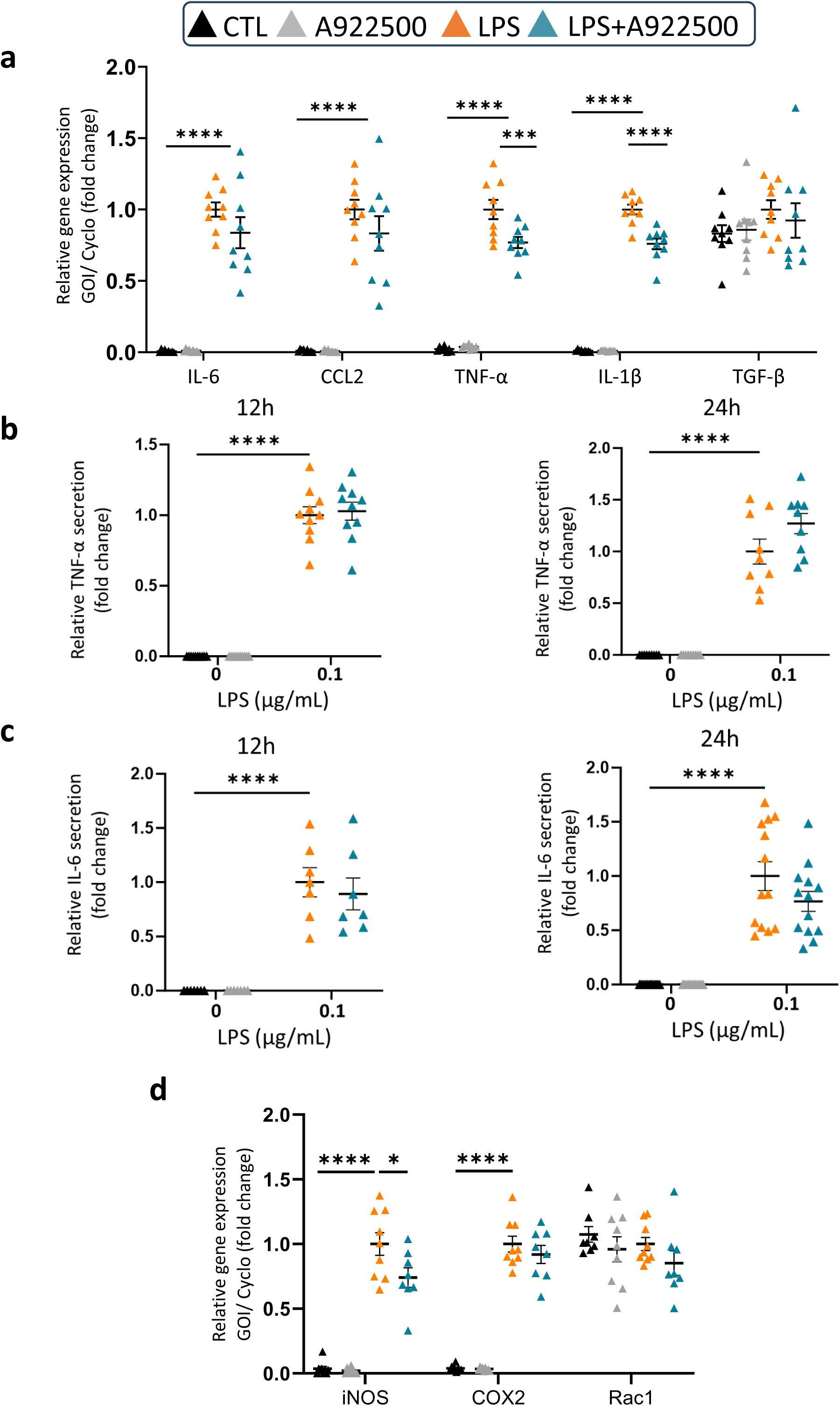
DGAT1 inhibition modulates microglial pro-inflammatory cytokine expression without altering cytokine secretion. Effect of treatment with 0.1 µg/mL LPS ± 500 nM A922500 on cytokine expression and secretion. **a)** Relative gene expression of cytokines in neonatal microglia measured by RT-qPCR (n = 8-9). Relative extracellular cytokine secretion at 12 h and 24 h of **b)** TNF-α and **c)** IL-6 (n=7-13). **d)** Relative gene expression of iNOS, COX2 and Rac1 was measured by RT-qPCR (n = 8-9) in microglia treated 6 h with 0.1 µg/mL LPS ± 500nM A922500. Two-way ANOVA with post-hoc Šídák, *p < 0.05, ***p < 0.001 **** p < 0.0001. IL-6; Interleukin-6, TNF-α; Tumor Necrosis Factor-alpha.

Furthermore, to assess if the decrease in cytokine expression translated into changes in cytokine secretion, ELISA were performed on culture supernatants collected after 12 h and 24 h of treatment with A922500 and/or LPS. The secretion of TNF-α and IL-6 induced by LPS was not significantly changed at either time points (**Fig.4b-c**). Altogether, these data suggest that DGAT1-dependent LD formation modulates the expression of specific cytokines without significantly affecting cytokine secretion.

To evaluate whether DGAT1 inhibition modulates LPS-induced cytokine expression through changes in reactive oxygen and nitrogen species, we measured the expression of iNOS (which converts L-arginine into nitric oxide during microglia activation), Rac1 (an activator of the NADPH oxidase complex involved in reactive oxygen species (ROS) production), and COX-2, which is typically upregulated in response to LPS-induced ROS^28^. As expected, treatment with 6 h of LPS increased expression of iNOS and COX-2 (**Fig.4d**). While Rac1 and COX-2 expression remained unchanged, DGAT1 inhibition significantly reduced LPS-induced iNOS expression by 26% (**Fig.4d**). This suggests that DGAT1 inhibition in microglia may selectively suppress nitric oxide–mediated inflammatory signaling, while not affecting ROS-associated pathways.

## DISCUSSION

In this study, we reveal the role of microglial DGAT1-mediated LD formation on inflammatory responses. Our data shows that acute inflammatory stimulation by LPS downregulates key enzymes involved in TG synthesis in primary neonatal microglia. Interestingly, while the canonical esterification pathway is downregulated, LPS simultaneously upregulates TMEM68, an enzyme involved in alternative TG and LD synthesis. Functional analysis demonstrated that microglial DGAT1 is the main enzyme responsible of LD formation in both oleate- and/or LPS-stimulated conditions, as its pharmacological inhibition with A922500 markedly reduced LD accumulation under both conditions. Inhibition of DGAT1 led to a decrease in LPS-induced expression of pro-inflammatory cytokines TNF-α and IL-1β, but had no effect on secretion. Together, we show that DGAT1-dependent LD formation is an important regulator of microglial inflammatory responses.

Consistent with previous studies, our findings show LD accumulation in microglia relies on DGAT activity ^15,22,29^. However, most studies examining the role of LD synthesis on microglial activity focused on disease-associated microglia from mouse models or iPSC derived from disease-associated microglia, where most changes in LD are dependent on DGAT2 ^22,30^. In contrast, our data reveal that DGAT1 is the main isoform expressed (no detectable expression of DGAT2) and that LD formation relies predominantly on DGAT1 activity in mouse microglia culture. The difference between studies may be related to the cell models and/or the inflammatory context.

As reported by us and others, treatment of primary microglia with LPS induced LD accumulation^11–13^. We found that LPS-induced LD formation was dramatically reduced by DGAT1 inhibition. Despite this marked reduction, DGAT1 inhibition did not affect the number of LD in unstimulated conditions. We previously reported that ATGL inhibition under basal conditions in primary microglia is sufficient to increase LD number^12^, suggesting that lipolysis is active in normal culture conditions and that the maintenance of the microglial LD pool may result from a balance between LD formation and hydrolysis. The fact that we did not observe changes in LD number in the presence of the DGAT1 inhibitor therefore indicates that basal LD formation may not be dependent on DGAT1.

This suggests the presence of a compensatory or alternative pathway for neutral lipid storage in microglial LD. Although DGAT2 expression was undetectable in our primary cultures, this does not necessarily exclude its contribution, as DGAT2 may be expressed below detection threshold. A compensatory effect of DGAT2 has already been demonstrated in adipocytes, where DGAT2 activity can sustain a low level of TG synthesis when DGAT1 is deleted^31^. In parallel, we found that TMEM68 was detectable and induced by LPS, while enzymes of the canonical TG-synthesis pathway were downregulated, highlighting a non-canonical esterification mechanism that may help maintain the pool of LD even when DGAT1 is inhibited. Taken together, these data suggest that, although DGAT1 is the primary driver of acute LD formation in response to oleate or LPS in primary microglia, additional pathways may contribute to maintaining a basal pool of microglial LD. Additional work will be needed to decipher the functional role of this alternative pathway on microglial activation.

We observed that LPS reduced the expression of the main TG esterification enzymes including DGAT1 while LPS promoted LD accumulation. This sounds counterintuitive but could be related to the fact that LPS also reduced ATGL mRNA and protein level in primary microglia^12^. Decreased ATGL activity and lipolysis may thus promote LD accumulation. The reduced expression of DGAT1 induced by LPS may help limit LD accumulation. This suggests that the pool of LD is dynamically coordinated by both the esterification and lipolytic arms to regulate inflammatory responses to LPS. Further studies will be needed to assess the respective contribution of esterification and lipolytic enzymes, including TMEM68, in LPS-induced LD accumulation and inflammation.

Despite a reduction of pro-inflammatory cytokines expression, blocking the esterification arm of LD did not affect cytokine secretion and did not reproduce the robust inhibitory effect on microglial pro-inflammatory responses observed when the lipolytic arm was inhibited^12,13^. Indeed, we showed that inhibition or deletion of ATGL under acute pro-inflammatory conditions strongly reduces both the expression and secretion of pro-inflammatory cytokines^12^. This difference could be explained by the fact that in presence of the DGAT1 inhibitor, the residual pool of LD may be mobilized via lipolysis to generate fatty-acid derived signals that sustain microglial activation. These observations suggest that a complete suppression of LD formation through simultaneous inhibition of DGAT1/2 and/or TMEM68 might be required to reproduce the phenotype observed with ATGL inhibition.

We expected that blocking DGAT1 could have repercussions on microglial oxidative pathway, given the established role of LD in sequestering free fatty acid into TG to prevent cellular stress^32^. Surprisingly, DGAT1 inhibition in presence of LPS did not affect expression of Rac1 or COX-2, two markers associated with ROS production in microglia. However, iNOS expression was decreased by DGAT1 inhibition in LPS-treated microglia, suggesting that DGAT1 may regulate nitric oxide–mediated inflammatory signaling.

Taken together, our findings identify DGAT1 as the primary enzyme responsible for LD formation and shaping acute inflammatory responses in neonatal microglia. This coordination between LD synthesis and breakdown opens new perspectives on how the dynamics of TG metabolism regulate microglial activation. This may open the way for new strategies to modulate microglial activity and control neuroinflammation in physiopathological contexts.

## METHODS

### 2.1. Animal ethics

Animal studies were conducted in accordance with guidelines of the Canadian Council on Animal Care (protocol # CM23041TAs). C57BL6/J mice purchased from Jackson Laboratories were used to generate pups for primary microglia. Mice were group housed on a 12 h dark-light cycle at 22–23 °C, with *ad libitum* access to standard irradiated chow diet (Teklad) and water.

### 2.2. Cell culture

#### 2.2.1. Primary neonatal mouse microglia

Primary neonatal mouse microglia were isolated from the forebrain of P1-2 mice (males and females pooled) using a method we described previously ^12,33^. Briefly, cerebellum, olfactory bulbs, and meninges were removed before rough homogenization with a blade. The homogenate was centrifuged at 500 g for 1 min and supernatant discarded. Homogenate was digested for 15 min in enzymatic solution (0.2 µm filter sterilised 4.5 U/mL papain [Worthington Biochemical, #LS003120], glucose (Fisher, BP350500, 5 mg/ml), cysteine (Fisher, 0.2 mg/ml), 1x DNAse [Worthington Biochemical, #LS006333], in PBS) at 37 °C. The digest was centrifuged at 500 g for 2 min and supernatant discarded. The digest was mechanically disrupted with a serological pipette in complete media (DMEM [Gibco, 11965–092] 4.5 g/dL glucose, 10 % FBS [Gibco, 10437-028], 50 U/mL penicillin, 50 µg/mL streptomycin [Gibco, 15070-063]). Cell suspension was filtered through a 70 µm cell strainer, centrifuged at 500 g for 5 min and resuspended in complete media. Cell suspension was plated in T75 flasks (50 % forebrain/flask) and allowed to adhere for 7 days. Media was changed when astrocyte monolayer reached confluency. When microglial cells began to detach from the astrocyte layer after another 5 days in culture, microglia were dislodged by tapping the side of the flask and media containing microglia was centrifuged at 500 g for 10 min at RTP. Cells were counted using a hemocytometer and seeded at an appropriate density. Complete media was added to the experimental dishes and microglia were allowed to adhere for 24 to 48 h prior to the start of the experiment.

#### 2.2.2. Cell culture treatments

The culture media was replaced 2 h prior to treatment with medium (DMEM [Winsent, 319–061-CL] 5 mM glucose) containing 0.25 mM oleic acid (Nu-chek Prep, #S-1120) pre-complexed to 0.27% Bovine Serum Albumin (BSA, Multicell, #800-095-EG) and/or 500 nM A922500 (Sigma, A1737-1MG). At time of treatment, medium was replaced with medium containing A922500 (500 nM), or vehicle (0.1 % v/v, DMSO, Sigma, D2438-5x10ml) with or without LPS (0.1 µg/mL, O127:B8, Sigma, L4516-1MG) or 0.25 mM oleic acid.

#### 2.2.3. Cell viability and cytotoxicity

The ApoTox-Glo kit was used to measure cell viability and cytotoxicity after 6 h treatment according to the manufacturer’s directions (Promega, G6320). Cells were treated with 0.1 µg/ml LPS and A922500 (500 nM) or vehicle (0.1 % v/v DMSO) for 6 h.

### 2.3. Lipid droplet imaging

#### 2.3.1 Confocal microscopy imaging

Primary microglia derived from pups were plated at a density of 1.5×10^5^ on 13 mm diameter glass coverslips in 24 well plates. After 6 h treatment, cells were fixed for 20 min with 4 % paraformaldehyde at 37 °C. Intracellular LD were visualized using the neutral lipid stain Lipidspot^488^ (10 µM, DMSO, Fisher, NC1669425) for 2 h. 1:1500 Hoescht ^33342^ (Invitrogen, #H3570) was used to identify nuclei. After staining, cells were mounted in ProLong Gold antifade mountant (Invitrogen, P36934) and allowed to set prior to imaging (64x oil-immersion lens, Confocal Leica Stellaris 8). LD were manually counted in ImageJ, with a minimum of 85 cells/coverslip (FIJI; ^34^).

#### 2.3.2 High throughput imaging

Primary microglia derived from pups were plated at a density of 4.0 x10^4^ on 96-well plates (PerkinElmer, #6055302). Cells were treated with DMEM (Multicell, #319-060-CL), 5 mM glucose and 0.25 mM oleate (Nu-chek Prep, #S-1120) pre-complexed to 0.27% BSA (Multicell, #800-095-EG), 500 nM A922500, 0.1 µg/ml LPS or 0.1% DMSO for 6 h as described in the figure legends. Cells were fixed using 4% paraformaldehyde for 15 min and neutral lipids were stained using 1:500 5 mM Bodipy ^493/503^ (Sigma, #D3922) while nuclei were stained using 1:1500 Hoechst ^33342^. Plates were imaged using Operetta High Throughput screening system and analyzed using Harmony High-Content Imaging and Analysis Software. For analysis, one experimental N consisted of the average of 6 wells in which, in every well, 9 areas were randomly selected for quantification.

### 2.4. RNA analyses

#### 2.4.1. Cell collection for RNA extraction

Primary mouse microglia were plated at a density of 3.0×10^5^ in 6 well plates and treated for 6 h (section 2.2.2) prior to RNA extraction with Trizol (Invitrogen, 15596-018) as we described^12,33^

#### 2.4.2. RNA extraction, cDNA synthesis and RT-qPCR

Extracted RNA was checked for purity and concentration using the Nanodrop 2000. For RNA extracted from neonatal microglia or tissue, cDNA was synthesised from 1 µg of RNA with M−MuLV reverse transcriptase (Invitrogen, 28025-013) and random hexamers (Cedarlane, 26-4000-10) as specified by the manufacturer. cDNA was diluted 1:10 prior to qPCR. qPCR master mix was composed of QuantiFast SYBR Green PCR kit (Qiagen, 204076) and forward and reverse primers (1 µM) for the relevant gene of interest as we described^12^. Primer sequences used are provided in Table 1. qPCR was performed for 40 cycles using the Corbett Rotor-Gene 6000 (Qiagen) and quantified using the standard curve method using Rotorgene Q series software (v2.3.1). Genes of interest were normalised to 18S or Cyclophilin. Gene expression is represented as fold change from mean of LPS-treated control groups after normalisation.

**Table 1.**
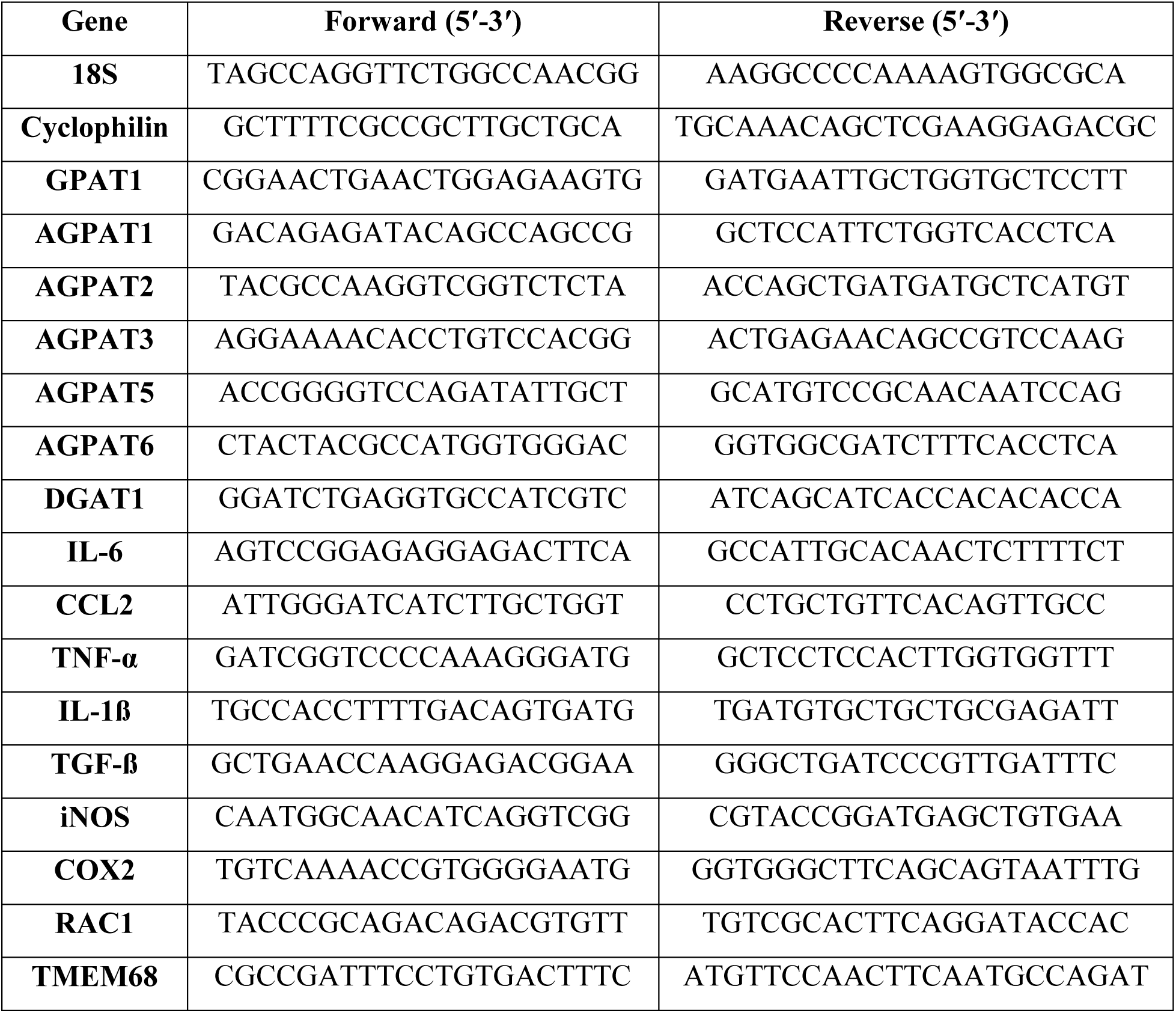
Primers sequence.

### 2.5. Cytokine secretion

Primary microglia derived from pups were seeded at a density of 4.0 x10^4^ in 96 well plates and treated for 12–24 h prior to collection of the extracellular media. TNF-α and IL-6 secretion into the extracellular media was quantified using DuoSet ELISAs (Bio-techne; MTA00B and DY406-05 respectively), as directed by the manufacturer and normalized to cell number as we described^12^. Cytokine secretion is represented as fold change from mean of LPS-treated control groups after normalisation.

### 2.6. Statistics

Data is presented as mean ± S.E.M. Data was prepared for analysis in Microsoft Excel and analysed and prepared for presentation using GraphPad Prism (v10.1.1). Intergroup comparisons were performed using Two-way ANOVAs with post-hoc Tukey’s or Šídák’s tests, One-way ANOVAs with post-hoc Tukey’s, Student’s t-test or Mann-Whitney as appropriate. Specific tests are detailed in figure legends. P<0.05 was considered significant.

## ACKNOWLEDGMENTS

We acknowledge the Cellular imaging and the Cell physiology core facilities at the CRCHUM. This work was supported by grants to T.A. from the National Natural Sciences and Engineering Research Council (NSERC, RGPIN-2025-04602) and the Canadian Institutes for Health Research (CIHR, PJT153035). J.L.R. was supported by a postdoctoral fellowship from Fonds de Recherche Québec-Santé (FRQS). F.B. was supported by scholarships from CRCHUM and Faculté de Médecine of Université de Montréal.

## AUTHOR CONTRIBUTIONS

F.B., D.M and J.L.R. performed cell culture, qPCR, ELISA, histology and imaging. D.R helped with mice breeding. F.B., D.M. J.L.R, and T.A. contributed to conceptualization, experimental design, data interpretation and manuscript revisions. F.B. and T.A. wrote the manuscript.

## ETHICS APPROVAL

All animal studies were conducted in accordance with guidelines of the Canadian Council on Animal Care and the Institutional Animal Care Committee of CRCHUM (protocol # CM23041TAs).

