## Supplemental Figures 1 and 2 for "DGAT1-dependent lipid droplet synthesis in microglia attenuates neuroinflammatory responses to lipopolysaccharides"

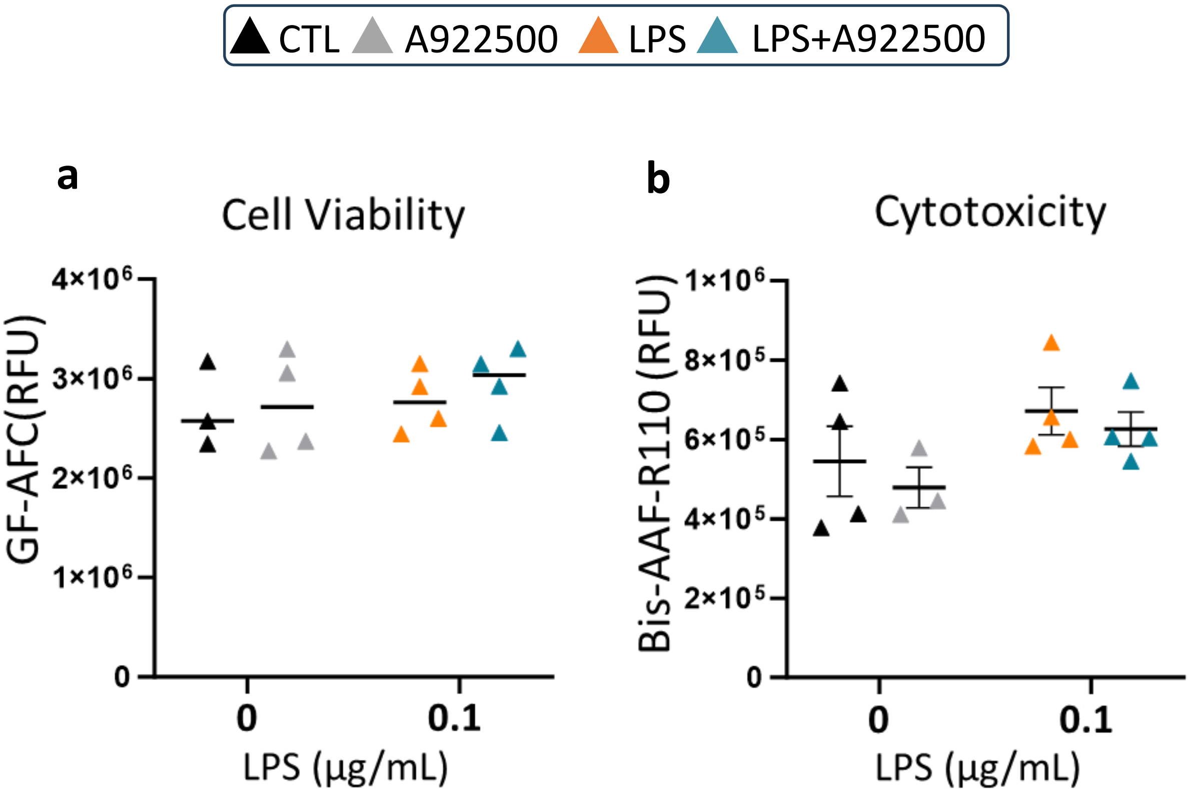


**Figure S1. Treatment of primary microglia with LPS or DGAT1 inhibitor A922500 does not impact microglia cell viability and cytotoxicity.** **a)** Cell viability and **b)** cytotoxicity of primary neonatal microglia treated with 6 h of 0.1 µg/mL LPS ± 500 nM A922500 (n = 3-4). Two-way ANOVA with post-hoc Šídák

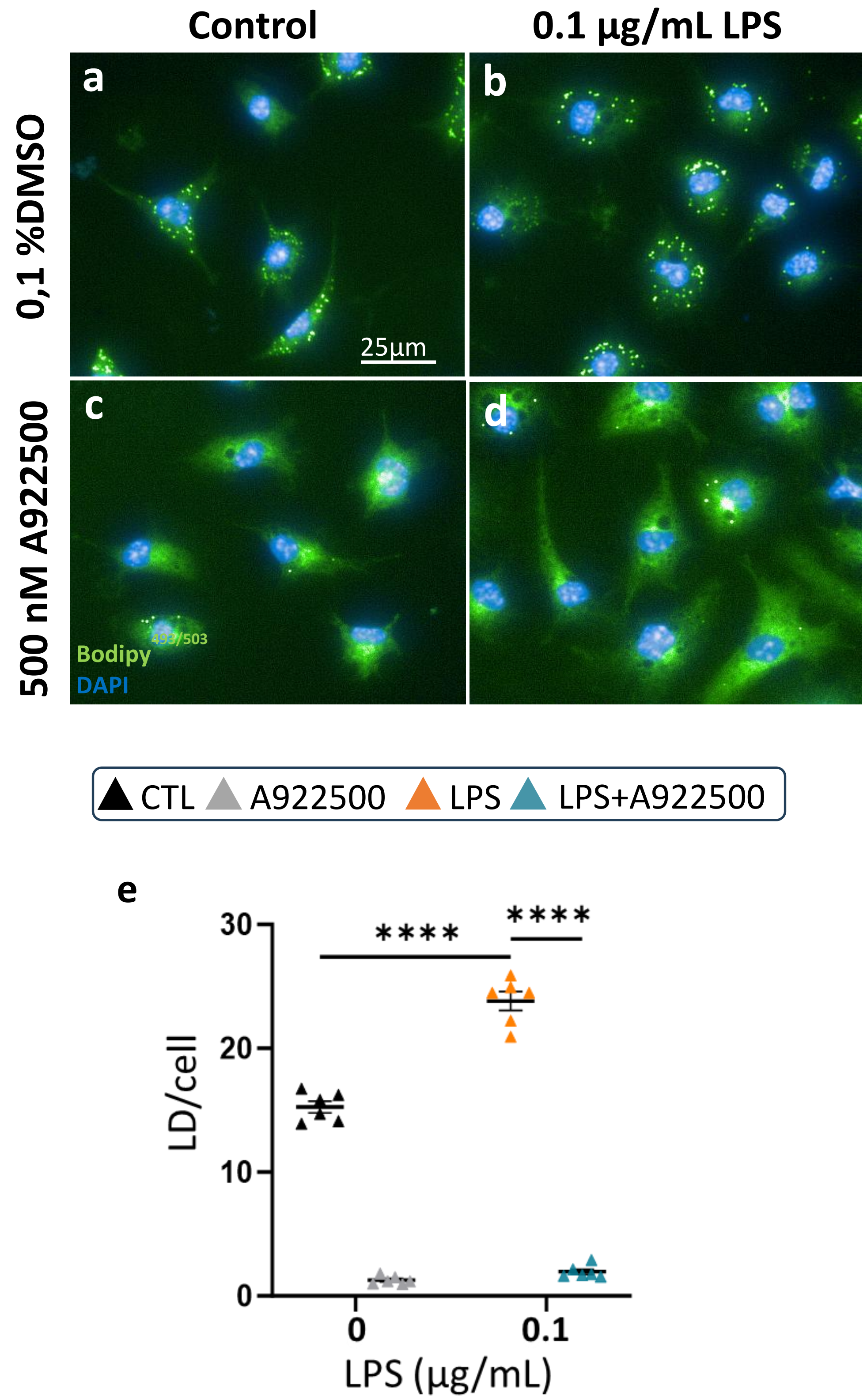
Control
0.1 µg/mL LPS
0,1 %DMSO
500 nM A922500
a
b
c
d
25µm
Bodipy 493/503
DAPI
CTL
A922500
LPS
LPS+A922500
e
LD/cell
LPS (µg/mL)
****
****
30
20
10
0
0
0.1

**Figure S2. DGAT1 is required for oleate-induced LD synthesis in primary microglia under pro-inflammatory conditions.** Effect of treatment with 0.25 mM oleate pre-complexed with 0.3% BSA, with or without 500 nM A922500 and/or 0.1 µg/mL LPS on LD number. **a-d)** Representative images of primary neonatal microglia pre-treated and maintained in oleate during the 6 h treatment with LPS or A922500, stained with BODIPY (green) and Hoechst (blue), 25 µm scale. **e)** Average number of LD per cell treated with LPS or A922500 (n =12, 9 fields of view/n, ~ 2700 cells counted/conditions). Two-way ANOVA with post-hoc Šídák, **** $p < 0.0001$.
